# Epigenetic inactivation of oncogenic brachyury (TBXT) by H3K27 histone demethylase controls chordoma cell survival

**DOI:** 10.1101/432005

**Authors:** Lucia Cottone, Edward S Hookway, Adam Cribbs, Graham Wells, Patrick Lombard, Lorena Ligammari, Anthony Tumber, Roberto Tirabosco, Fernanda Amary, Tamas Szommer, Catrine Johannson, Paul E Brennan, Nischalan Pillay, Udo Oppermann, Adrienne M Flanagan

**Affiliations:** UCL Cancer Institute, University College London Cancer Institute, London, WC1E 6BT, UK; Botnar Research Centre, Nuffield Department of Orthopaedics, Rheumatology and Musculoskeletal Sciences, University of Oxford, Oxford, OX3 7LD, UK; Structural Genomics Consortium, University of Oxford, OX3 7DQ, Oxford, UK; FRIAS - Freiburg Institute of Advanced Studies, 79104, Freiburg, Germany; Department of Histopathology, Royal National Orthopaedic Hospital, Stanmore, Middlesex HA7 4LP, UK

## Abstract

The expression of the transcription factor *brachyury* (*TBXT*) is normally restricted to embryonic development and its silencing after mesoderm development is epigenetically regulated. In chordoma, a rare tumour of notochordal differentiation, TBXT acts as a putative oncogene, and we hypothesised that its expression could be controlled through epigenetic inhibition. Screening of five chordoma cell lines revealed that only inhibitors of the histone 3 lysine 27 demethylases KDM6A (UTX) and KDM6B (Jmjd3) reduce *TBXT* expression and lead to cell death, findings validated in primary patient-derived culture systems. Pharmacological inhibition of KDM6 demethylases leads to genome-wide increases in repressive H3K27me3 marks, accompanied by significantly reduced TBXT expression, an effect that is phenocopied by the dual genetic inactivation of *KDM6A/B* using CRISPR/Cas9. Transcriptional profiles in response to a novel KDM6A/B inhibitor, KDOBA67, revealed downregulation of critical genes and transcription factor networks for chordoma survival pathways, whereas upregulated pathways were dominated by stress, cell cycle and pro-apoptotic response pathways.

This study supports previous data showing that the function of TBXT is essential for maintaining notochord cell fate and function and provides further evidence that TBXT is an oncogenic driver in chordoma. Moreover, the data suggest that TBXT can potentially be targeted therapeutically by modulating epigenetic control mechanisms such as H3K27 demethylases.

## Introduction

Normal embryonic development requires a finely co-ordinated process of temporal and spatial gene expression which is controlled through epigenetic mechanisms (1, 2). Prenatal dysregulated expression of key developmental genes can lead to specific clinical phenotypes, such as MLH1 and FMR1 repression in hereditary non-polyposis colon cancer and in fragile X syndrome respectively (3). In addition, aberrant epigenetic mechanisms also have been implicated in the pathogenesis of a diverse range of malignancies (4).

During vertebrate development, mesoderm specification is tightly regulated by the prototypical T-box transcription factor T (TBXT, also known as T, and brachyury) which through exquisitely orchestrated caudo-cranial morphogenic gradients, controlled through FGF and Activin signalling, ultimately transforms the nascent dorsal mesoderm into the rudimentary axial skeleton, the notochord (5–7). *TBXT* is silenced in the human fetus at approximately 12 weeks and consequentially the notochord recedes prenatally. In embryonic stem cells and mesoderm-derived differentiating adipogenic cells, regulation of *TBXT* gene expression is achieved through epigenetic regulatory mechanisms. These include the action of lysine demethylases, in particular lysine demethylase 6A (KDM6A, also known as UTX) and lysine demethylase 6B (KDM6B, JMJD3) (hereafter referred to as KDM6A/B) both demethylases of tri-methylated histone 3 lysine 27 residues (H3K27me3) (8, 9), the recruitment of histone deacetylase 1 (HDAC1) (10) and CpG island methylation (11, 12).

In adult tissue aberrant *TBXT* expression is seen in chordoma, a rare cancer of the axial skeleton showing notochordal differentiation (13, 14), in the benign notochordal cell tumour, - the precursor of chordoma, and in the haemangioblastoma a rare tumour reminiscent of the embryonic hemangioblast. Individuals diagnosed with chordoma have a median of 7 years survival, and they have seen little benefit from genomic studies seeking tractable therapeutic targets such as protein kinases (15). Small molecule-focused compound screens against chordoma cell lines *in vitro* have also yielded limited success with few compounds showing inhibition of cell growth: EGFR inhibitors were one of the rare exceptions, results which have led to a phase 2 clinical trial (16, 17). Although encouraging, this treatment is unlikely to benefit all patients as not all cell lines were responsive, and the rapid development of resistance to EGFR inhibitors in clinical practice is widely documented (18).

TBXT plays a pivotal role in the development of chordoma (19). Specifically, *TBXT* expression is seen in chordoma but is silenced in other tumours of mesodermal origin (20). At a genomic level germline tandem duplication of *TBXT* is a key genetic predisposition event in familial chordoma (21), and 97% of patients with sporadic chordoma harbour the rs2305089 SNP within the DNA binding domain of *TBXT* conferring a significantly increased risk of developing the disease (22). Furthermore, at a functional level, *TBXT* acts as a master regulator of an elaborate oncogenic transcriptional network (19) and its silencing in chordoma cell lines *in vitro* causes growth arrest (23, 24). Despite this substantial body of evidence demonstrating that TBXT expression confers oncogenic properties in chordoma, the key mechanisms for regulating its expression have not been reported. This cannot be accounted for by the somatic copy number gain of *TBXT* as this is only observed in up to 27% of the sporadic cases (15). In this study we set out to investigate the epigenetic regulation of *TBXT* in chordoma using a focused library of mechanistically defined inhibitor tool compounds with the aim of identifying vulnerable signalling pathways, which could be exploited for the development of novel treatments for this disease.

## Results

### H3K27 demethylases regulate cell viability in chordoma

To identify epigenetic processes of biological importance in chordoma, we screened a library of tool compounds (**Dataset S1**) (25) targeting proteins involved in chromatin biology including several components of epigenetic readers, writers and erasers of a ‘histone’ or ‘chromatin code’, against five well characterised chordoma cell lines (www.chordomafoundation.com). A resazurin-based viability assay was used as the primary anti-proliferative readout (**Fig. 1A**). Decreased cell viability was observed in all cell lines in response to HDAC inhibitors as previously reported (26). Broad-spectrum inhibitors of the Jmj-type of lysine demethylases were seen to reduce cell viability, as was the more specific inhibitor of KDM6A/B GSK-J4 (27) (**Fig. 1A-B**) but not the regio-isomer control compound GSK-J5, an observation that led us to focus or investigations on the inhibitors of H3K27 demethylases. Decreased viability was also observed with KDOBA67, a novel cell-permeable hydroxyl derivative of GSK-J4 that does not require activation by intracellular esterases (**Fig. 1A-B and Fig. S1)**. KDOBA67 displayed *in vitro* inhibitory activity in the low micromolar range with IC50 values of 2–5 μM in various chordoma cell lines (**Fig. 1C**), and led to proliferation arrest and induction of apoptosis **(Fig. 1D-G and Fig. S2**). As expected, the regio-isomer control compound GSK-J5 did not display anti-proliferative effects (**Fig. 1A-G and Fig. S2**).

**Figure 1.**
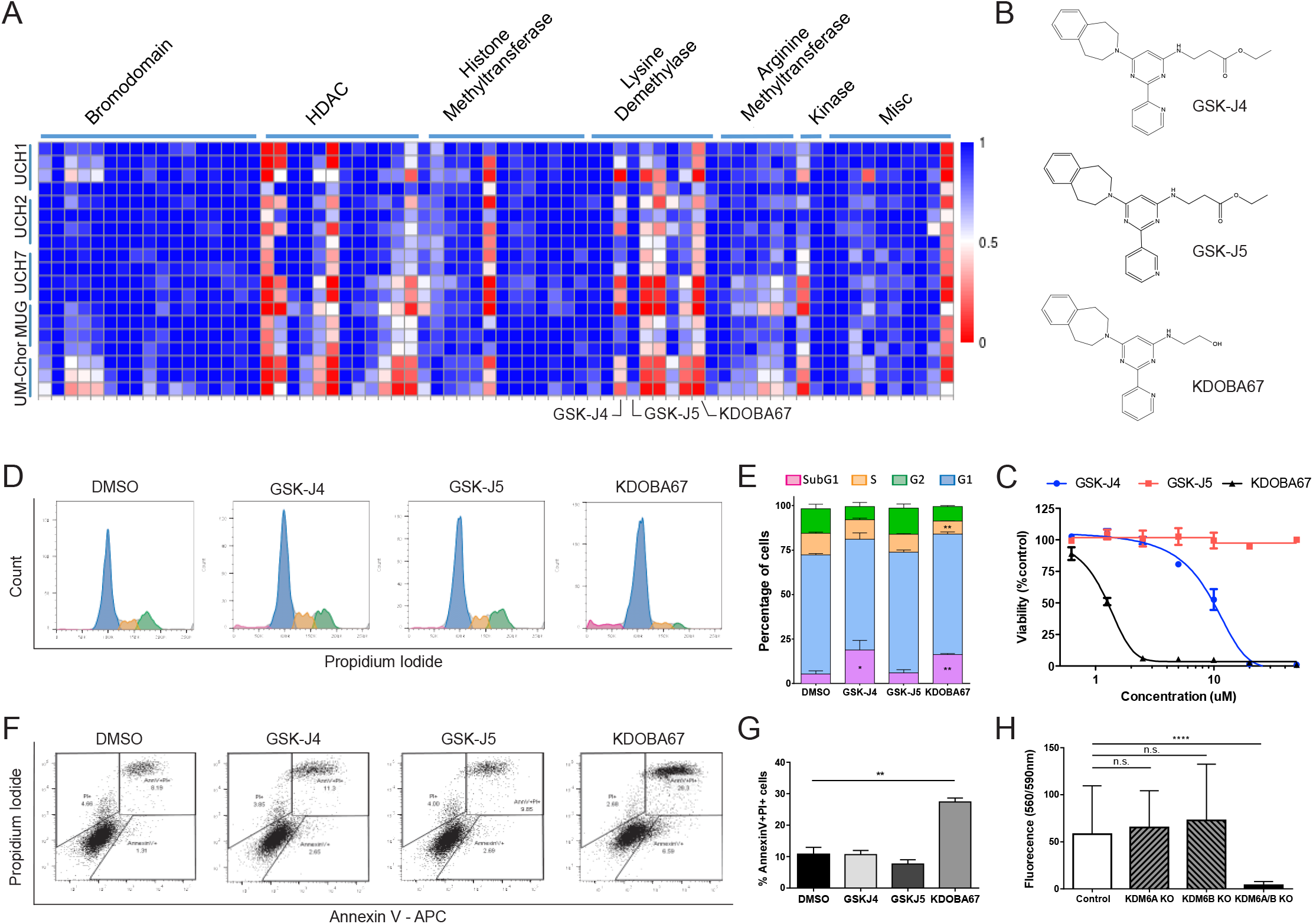
Focused epigenetic library screening identifies H3K27 lysine demethylases as potential therapeutic target in chordoma. (**A**) Screening of 90 small molecule probes with validated activity against enzymes involved in chromatin biology on 5 chordoma cell lines. Each row represents a separate replicate and each column an individual compound. Columns have been grouped together by inhibitor class (c.f. also to **Dataset S1**). Values plotted as fractional viability compared to vehicle (DMSO) control. (**B**) Molecular structures of GSK-J4, GSK-J5 and KDOBA67. (**C**) Dose response curves for GSK-J4, GSK-J5 and KDOBA67 in UM-Chor. (**D-G**) Cell cycle changes (**D-E**) and cell death analysis assessed by AnnexinV-PI staining (**F-G**) in response to the compounds after 72 hours of treatment. Representative histogram/dot plots and quantification of 3 independent experiments, with 3 replicates per condition (c.f. also **Fig. S2)**. (**H**) Viability measured by Alamar blue of UCH1 following CRISPR/Cas9 editing of *KDM6A/B* or both. *p ≤0.05, **p ≤0.01, ***p ≤0.001, ****p ≤0.0001.

To confirm that the observed inhibitory growth effects with the tool compounds were due to on-target inhibition of KDM6A/B, we used the CRIPSR/Cas9 system to introduce non-sense mutations in both *KDM6A* and *KDM6B* genes (**Fig. S3A-B**). Targeting either gene alone was not sufficient to reduce cell viability but cells in which both genes had been edited showed a significant (p-value < 0.0001) decrease in viability and proliferation (**Fig. 1H and Fig. S3C-G**). The requirement for inactivation of both *KDM6A/B* is supported by the finding that the cell line MUG-Chor which contains a biallelic deletion of *KDM6B* only required CRISPR/Cas9 inactivation of *KDM6A* (**Fig. S3F-H**).

### *TBXT* expression is regulated by H3K27 demethylases

Given the central role of TBXT in chordoma biology, we next asked whether inhibition of KDM6A/B would effect expression of *TBXT* in chordoma cell lines. Western blot and qPCR analysis confirmed that treatment with either GSK-J4 or KDOBA67, but not with the control compound GSK-J5, reduced TBXT in all chordoma cell lines tested (**Fig. 2A-B and Fig. S3 J-K)**. Reduction in TBXT expression was not only seen in cell lines containing a diploid copy number of the gene (UCH1 and UM-Chor) but also in the MUG-Chor cell line that contains multiple copies of the *TBXT* gene (www.chordomafoundation.org). The effect of KDOBA67 was substantiated further by demonstrating both anti-proliferative effects (**Fig. 2C**) and decreased TBXT expression (**Fig. 2D**) on treatment of primary cultures of patient-derived chordoma cells. The reduction of *TBXT* expression was reproduced by inactivating genetically KDM6A/B in chordoma cell lines using CRISPR/Cas9 (**Fig. 2E** and **Fig. S3I**).

**Figure 2.**
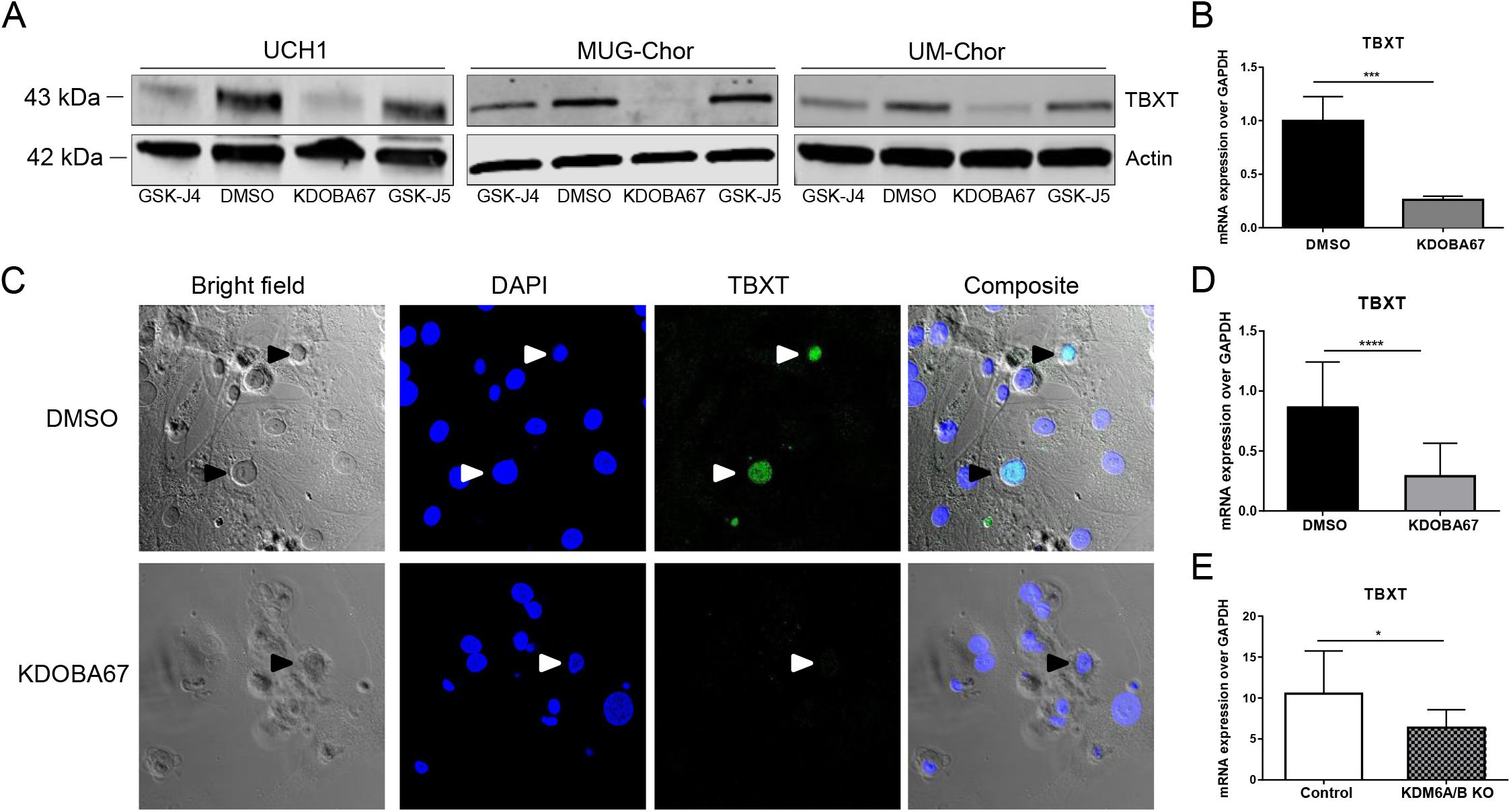
Specific inhibition of H3K27 lysine demethylase leads to inactivation of TBXT in chordoma. (**A**) Protein expression of TBXT in chordoma cell lines, following treatment with KDOBA67 as assessed by western blotting. Beta actin is used as an endogenous control. (**B**) TBXT expression in UCH1 following KDOBA67 treatment as assessed by qPCR. (**C**) Cell death and reduction in expression of TBXT (harrow heads) as shown by immunofluorescence in primary patient-derived chordoma cultures treated with KDOBA67 or control DMSO. Chordoma cells identified by the expression of TBXT are interspersed in tumour-derived stromal cells. TBXT/green, DAPI/blue. 40 X magnification. (**D**) *TBXT* transcript level is reduced in primary chordoma cultures upon treatment with KDOBA67 as assessed by qPCR. N=4 primary tumours, at least 3 replicates per condition. (**E**) Expression of *TBXT* is reduced in UCH1 upon double KO of *KDM6A/B*, assessed by qPCR. Quantification of 2 independent experiments, with 2 replicates per condition. *p ≤0.05, **p ≤0.01, ***p ≤0.001, ****p ≤0.0001.

### Inhibition of KDM6A/B alters the chromatin state at the TBXT locus

To investigate whether the inhibition of KDM6A/B regulates *TBXT* epigenetically, and reflects the effect of ablation of these genes in development (8, 9), we examined the distribution of histone marks and histone variants surrounding the *TBXT* locus in chordoma cell lines. ChIP-PCR demonstrated enrichment of H3K4me3 and H3.3, chromatin marks associated with active gene transcription, at the *TBXT* promoter in 3 chordoma cell lines known to express *TBXT* but not in the osteosarcoma cell line U2OS that does not express *TBXT* **(Fig. 3A-B)**. In contrast, the repressive marks H3K27me3 and H3K9me2 were enriched in the osteosarcoma cell line but not in the chordoma cells (**Fig. 3C-D**). Examination of DNA methylation at the *TBXT* locus showed consistent low methylation in the promoter region and at the transcription start site not only in primary chordoma tumours (n=35) but also in a range of primary bone tumours and primary solid tumours that are known not to express *TBXT* (**Fig. 3E and Fig. S4**). These data imply that alteration in promoter DNA methylation does not play a significant role in the regulation of *TBXT* in chordoma and a range of malignancies.

**Figure 3.**
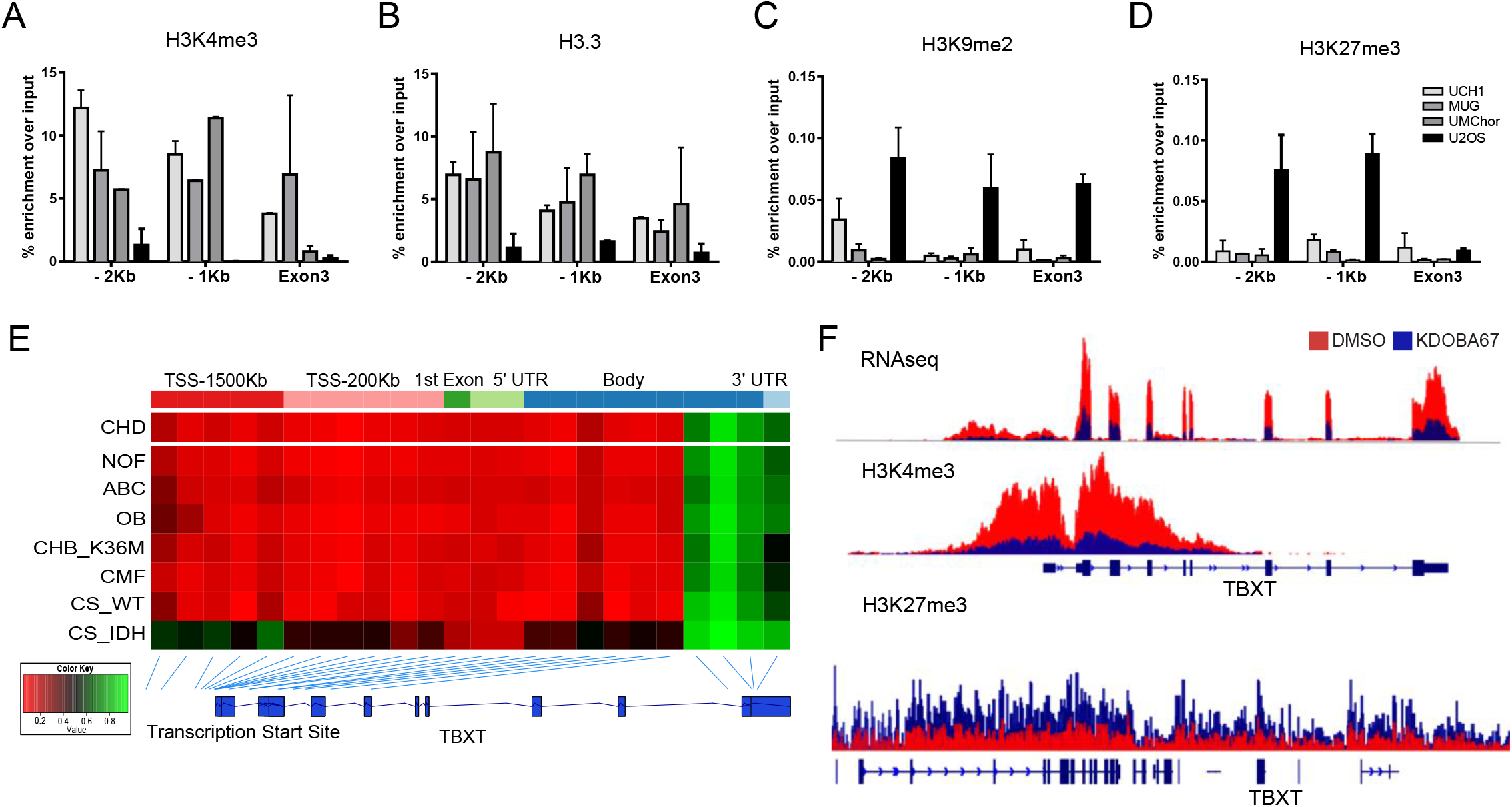
Epigenomic and transcriptomic profiling in response to H3K27 lysine demethylase inhibitors in chordoma cell lines. ChIP-PCR of H3K4me3 (**A**), H3.3 (**B**), H3K9me3 (**C**) and H3K27me3 (**D**) in 3 chordoma cell lines (UCH1, MUG-Chor1, UM-Chor1) and the osteosarcoma cell line U2OS. (**E**) The promoter region of *TBXT* is hypomethylated in all primary human chordomas (CHD, n=35) and in other mesenchymal tumours not associated with the expression of TBXT. From top to bottom: non-ossifying fibroma (NOF, n=12), aneurysmal bone cysts (ABC, n=9), osteoblastoma (OB, n=12), chondroblastoma harbouring H3.3-K36M mutation (CHB-K36M, n=17), chondromyxoid fibroma (CMF, n=25), chondrosarcoma WT (CS-WT, n=2), chondrosarcoma-harbouring an IDH mutation (CHS-IDH, n=3). The average beta value for each probe in each cancer type is plotted with the position of the probe shown relative to the TBXT gene body. (**F**) RNA expression (top panel), H3K4me3 coverage (middle panel) and H3K27me3 coverage (bottom panel) for KDOBA67 treated (blue) and DMSO-treated (red) cells. A decrease in the level of H3K4me3 around the *TBXT* transcription site was observed with a concurrent increase in the level of H3K27me3 over the gene locus. RNA expression is average for UM-Chor and MUG-Chor. ChIP signal is average for UCH1, UM-Chor and MUG-Chor.

To investigate further the chromatin landscape surrounding *TBXT* and the global changes induced by inhibition of KDM6A/B, we performed quantitative ChIP-seq of H3K27me3 and H3K4me3 marks. Treatment with KDOBA67 led to an increase in H3K27me3 and a decrease in H3K4me3 levels at the *TBXT* locus (**Fig. 3F**), and these findings were associated with significantly decreased expression of *TBXT* as assessed by RNAseq (**Fig. 3F** and **Dataset S2**; log2-fold changes −2.1 (MUG-Chor), −1.2 (UCH1), −0.96 (UM-Chor), −1.5 (UCH7) (p-value < 0.01)).

### H3K27me3 demethylase inhibitor treatment leads to genome-wide alterations in H3K4 and H3K27 methylation

Methylation of histone residues is a ubiquitous post-translational modification, and inhibition of a demethylase would be expected to have a broad response as opposed to a targeted effect on a single gene. We analysed the distribution of H3K4me3 and H3K27me3 throughout the genome at 48 hours after exposure to KDOBA67 and found there was global suppression of the level of H3K4me3 with a global increase in the level of H3K27me3 (**Fig. 4A-B**). Treatment with KDM6A/B inhibitors did not lead to a change in the location of H3K4me3 peaks but rather to a global decrease in levels of H3K4me3 (**Fig. 4B**). As reported previously for GSK-J4 (28) we also found that KDOBA67 exerts some inhibitory activity *in vitro* against KDM5 (JARID1) enzymes, the principal H3K4me3 demethylases (**Table S1**). The observed decrease in H3K4me3 levels in our cell-based experiments is in stark contrast to the reported effect of specific H3K4me3 demethylase inhibitors which lead to a global cellular increase in H3K4me3 (29, 30). This indicates that the phenotypic response observed in our experiments is the result of the inhibition of KDM6A/B and not an off-target effect against KDM5 enzymes.

**Figure 4.**
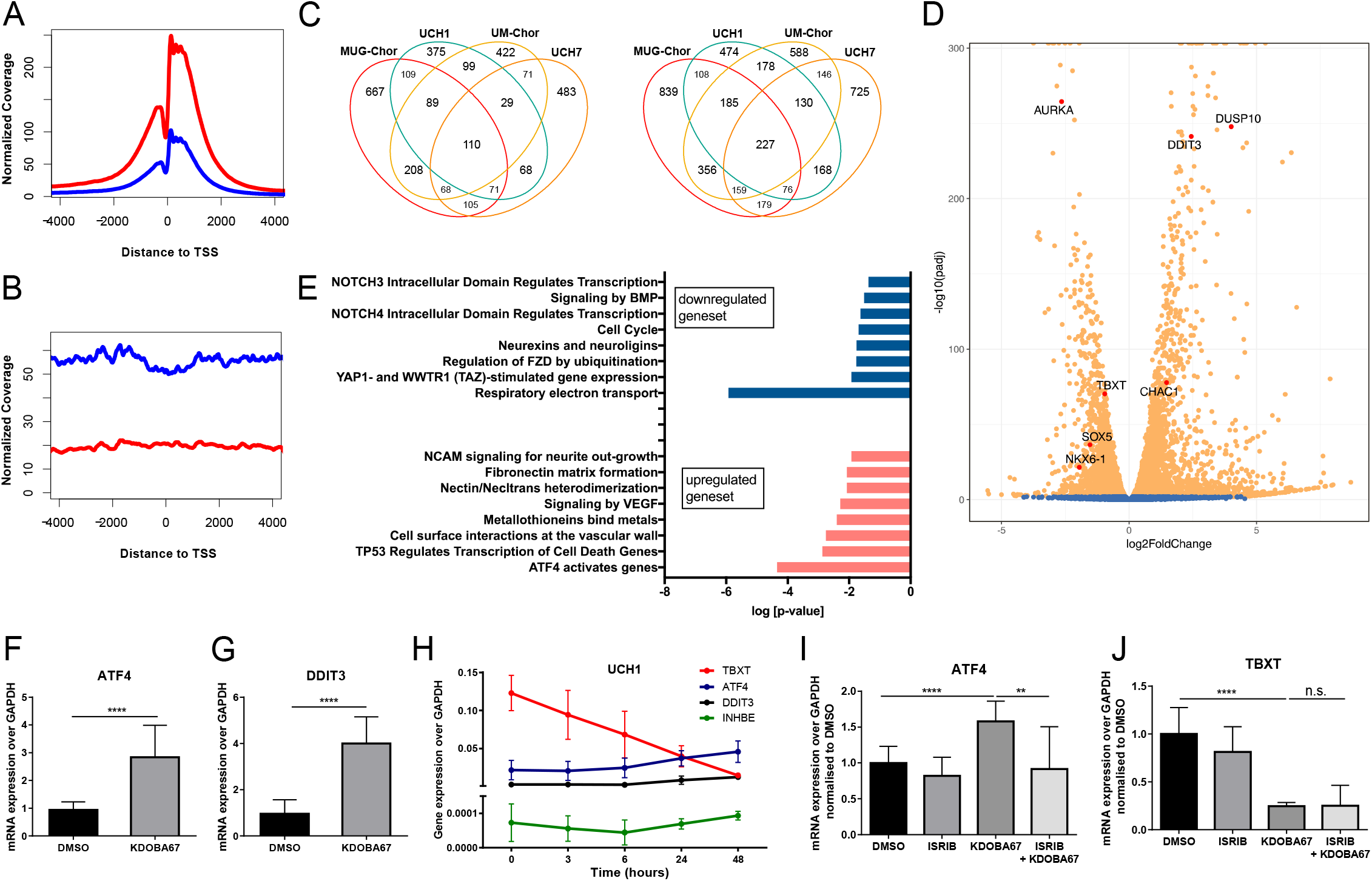
Epigenetic regulation of *TBXT* occurs independently of an ATF4 metabolic stress response. (**A-B**) Average H3K4me3 (**A**) and H3K27me3 (**B**) coverage across the transcription start sites (TSS) of all reference genes following 48 hours of DMSO and KDOBA67 (average for UCH1, UM-Chor and MUG-Chor). (**C**) Venn diagrams showing the numbers of downregulated (left) and upregulated (right) genes in 4 chordoma cell lines. (**D**) Volcano plot summarising differential expression in chordoma cells (UM-Chor) following treatment with KDOBA67 (c.f. also **Fig. S5** and **Dataset S2**). Blue, not significant; Orange, P-adjusted value <0.1. (**E**) Reactome analysis for UCH1, UCH7, UM-Chor and MUG-Chor in response to KDOBA67. (**F-G**) Expression of *ATF4* and *DDIT3* in primary chordoma cell cultures in response to KDOBA67, as assessed by qPCR on 4 primary chordoma samples. (**H**) *ATF4, DDIT3, INHBE* and *TBXT* regulation over a time course in response to KDOBA67 in UCH1 as assessed by qPCR. Results from 2 independent experiments, with 3 replicates per condition. Expression of *ATF4* (**I**) and *TBXT* (**J**) in UCH1 in response to treatment with KDOBA67 and ISRIB as assessed by qPCR. Quantification of 3 independent experiments, with 3 replicates per condition. *p ≤0.05, **p ≤0.01, ***p ≤0.001, ****p ≤0.0001.

### Inhibition of KDM6A/B reveals critical pathways for chordoma survival

We next investigated the implications of the global change in the chromatin landscape. The transcriptional profile of chordoma cell lines UCH1, UCH7, MUG-Chor and UM-Chor in response to KDOBA67 was interrogated by RNA sequencing at 48 hours after induction of inhibitor treatment (**Dataset S2 and Fig. S5A**) and this was found to result in significant perturbations in gene expression profiles (**Fig. 4C**). Within the downregulated gene sets *TBXT* and other transcription factors, such as the *SRY* box factor *SOX5* (31), a previously determined direct target of *TBXT* (19), and homeobox factors such as *NKX6–1* are identified **(Fig. 4D and Fig. S5 B-E**), suggesting a possible TBXT-controlled transcriptional network. Considering an overlapping set of 371 genes which are commonly regulated across all chordoma cell lines **(Fig. 4C)**, affected pathways are linked to chordoma biology (19) and related to cell movement, adhesion, extracellular matrix organisation, cell contact and signaling **(Fig. 4E)**. Importantly, this analysis reveals enrichment in categories such as apoptosis, mitochondrial impairment, cell cycle arrest and mitosis **(Fig. 4E)** with key genes such as *AURK, DUSP10* or the ATF4/*DDIT3*-target gene *CHAC1* being identified **(Fig. 4D and Fig. S5 B-E)**, findings which are consistent with the obtained anti-proliferative results **(Fig. 1)**.

Treatment of primary chordoma cells with KDOBA67 revealed a similar transcriptomic response to that seen in chordoma cell lines with increases in *ATF4/DDIT3*-target genes as measured by qPCR (**Fig. 4F-G and Fig. S6A-B**). The observation of an ATF4-driven stress response is in keeping with data observed in our previous studies and that of others, of the transcriptomic response to KDM6 inhibitors in human NK cells and neuroblastoma cells respectively (25, 32). As the ATF4-driven stress response leads to a global inhibition of translation as a mechanism for escaping diverse stressors (33), we set out to determine if the effect seen on expression of *TBXT* was a primary effect of KDM6 inhibition or if it occurred as a consequence of the induction of the stress response. Time course experiments looking at the expression of *TBXT* and the markers of the ATF4 response demonstrated that the expression of *TBXT* begins to decrease prior to the upregulation of genes associated with the ATF4 response (**Fig. 4H and Fig. S6C-D)**. To investigate this observation further, we treated chordoma cells with ISRIB (34), a compound that inhibits the induction of the ATF4 response by interfering with eIF2α phosphorylation: ISRIB did not prevent the downregulation of *TBXT* induced by KDOBA67 (**Fig. 4I-J and Fig. S6E-G**) but otherwise inhibited the ATF4 response. These data, in addition to the observation of the change in histone marks around the *TBXT* gene locus, demonstrate that the regulation of *TBXT* expression is directly mediated by KDM6 inhibition and does not occur secondary to the ATF4 response. The results obtained suggest that a network of transcription factors controlled by TBXT (19) is critical for cell survival, is regulated by H3K27 demethylases and can be epigenetically targeted to arrest chordoma cell proliferation and survival.

## Discussion

In this work we have identified that inhibition of H3K27 demethylases by KDOBA67, a derivative of the previously reported H3K27 demethylase inhibitor, GSK-J4, induces suppression of the embryonic transcription factor TBXT through a global increase of the repressive post-translational modification H3K27me3, followed by apoptosis of chordoma cells. The results highlight several important aspects in chordoma and epigenetic biology.

From a potential therapeutic standpoint, the number of small molecules that induce significant anti-proliferative effects in chordoma cells is limited (16). To our knowledge, this is the first time that such a compound has been shown consistently to induce cell death across a number of chordoma cell lines and to reduce simultaneously expression of *TBXT*. Previously, we and others have shown that treatment with EGFR inhibitors lead to decreased chordoma cell viability (16, 17, 35), however, the results are variable across different cell lines, and reduction of *TBXT* expression was seen in response to only one EGFR inhibitor, afatinib (16, 17). The finding that suppression of *TBXT* expression and reduced cell survival by H3K27 demethylase inhibition occurred in all tested chordoma cell lines and 4 genomically heterogeneous primary chordoma samples provides compelling evidence that the cellular response to the employed tool compound is consistent. Furthermore, the data imply that any chordoma expressing TBXT is likely to show similar responses to H3K27 demethylase inhibitors due to epigenetic deregulation of the *TBXT* locus. With TBXT being implicated in playing a fundamental role in the pathogenesis of chordoma, and apparently being expressed only by neoplastic cells in adult life, it appears to be an ideal therapeutic target (14, 15, 19, 21, 24, 36). Recognition of this is supported by the development of TBXT-specific poxviral vaccine not only for the treatment of chordoma but also for the minority of cases in which TBXT is expressed in more common cancers where it is supposedly involved in neoplastic epithelial-mesenchymal transition (37). The challenge in delivering a therapeutic agent is that transcription factors such as TBXT are often considered “undruggable” (38). However, our data show that it is feasible to block the function of *TBXT* in chordoma cells indirectly by decreasing its expression via manipulation of the chromatin environment. This strategy has been successfully used for targeting e.g. c-Myc with epigenetic BET bromodomain inhibitors such as JQ1 (39).

Understanding the mechanisms by which *TBXT* is expressed in chordoma is critical to understanding this disease. Previous work has implicated DNA promoter methylation in the expression programming of *TBXT* in development (40). However, our data show that the promoter region of *TBXT* shows low methylation not only in chordoma samples, but also in a range of sarcomas in which *TBXT* is not expressed, implying that low methylation of the *TBXT* promoter is not sufficient to drive gene expression. Our finding that KDM6A/B inhibition increases H3K27me3 levels, presumably leading to Polycomb Repressive Complex-mediated silencing of gene expression, provides further evidence that lysine demethylases are implicated in the physiological processes of silencing *TBXT* during embryonic development (8, 9).

Our study provides further insight into H3K27 demethylase functions in human tumour biology. KDM6/H3K27 demethylases possess in addition to their catalytic subunit additional domains that mediate protein-protein interactions (41). Since a non-enzymatic scaffolding function of H3K27 demethylases is necessary for developmental effects observed in mesoderm and vertebrate development (8, 9), and also to T-box transcription factor activity (42), it is of fundamental interest that inhibition of the enzymatic demethylase functions in the context of chordoma survival is sufficient to control tumour cell proliferation. Furthermore, the inhibitory effect on *TBXT* expression and associated cell death in response to *TBXT* gene silencing is only recapitulated by a concerted knockdown of both *KDM6A* and *KDM6B* genes. This significant overlap of chemical and genetic perturbations points to synergistic functions for both H3K27 demethylases in regulating the repressive chromatin environment in chordoma, and supports the cooperative function of KDM6A and KDM6B in maintaining the oncogenic expression of *TBXT* in chordoma.

In contrast to the cooperative function of KDM6A and KDM6B enzymes in chordoma, singular or even opposing roles of KDM6 enzymes have been noted in other malignancies. Whereas KDM6A is considered a tumour suppressor in many instances (43), KDM6B has been identified as oncogenic driver, e.g. in T-cell acute lymphoblastic leukaemia, controlling Notch1 and its target gene network (44). Knockdown experiments in chordoma cell lines have revealed that TBXT orchestrates a network of genes involved in the regulation of the cell cycle, in the production of extracellular matrix, growth factor and cytokine secretion, which are necessary for the survival of chordoma cells (19) and pertinent to notochord formation (16, 19), all of which are downregulated by KDOBA67 treatment. Novel in this scenario is the discovery of possible pro-apoptotic effects observed upon H3K27 demethylase inhibition leading to an ATF4- and DDIT3-driven stress response and highlighting this pathway as a possible therapeutic option in a range of different cancers (33).

In conclusion, our study provides evidence that H3K27 demethylases are major drivers in the maintenance of the oncogenic expression of *TBXT* in chordoma. In addition, we show that inactivation of H3K27 demethylases alters the expression of pathways critical to the survival of cancer cells, opening the road to further studies addressing this pathway as a potential molecular target.

## Materials and Methods

### Cell lines and primary cultures

5 well characterised human chordoma cell lines, U-CH1, U-CH2, U-CH7, MUG-Chor and UM-Chor (www.chordomafoundation.org), and the osteosarcoma cell line U2OS (ATCC, VA, USA) that lacked expression of TBXT and was used as a control, were cultured as described in **SI Materials and Methods**. Fresh sacral chordoma samples were obtained with consent from patients being treated at the Royal National Orthopaedic Hospital (RNOH), Stanmore, United Kingdom and processed as described in **SI Materials and Methods**.

### Screening of small molecule inhibitors, compound synthesis and compound treatments

To screen small molecules inhibitors, chordoma cell lines (U-CH1, U-CH2, U-CH7, MUG-Chor and UM-Chor) were treated with either a compound (a list of compounds and concentrations used is available in **Dataset S1**) or vehicle control (0.1 % DMSO) and the IC50 of lead compounds was calculated, all as described in **SI Materials and Methods**. In follow up experiments chordoma cell lines were treated with compounds at the IC80: GSK-J4 (5 μM, Cayman chemicals, An Arbor, USA), KDOBA67 (5 μM) and ISRIB (250 nM, Sigma Aldrich, St. Louis, MO, USA) for 48 hours (or for 72 hours in apoptosis and cell cycle experiments only). Primary chordoma cultures were treated with KDOBA67 (10 μM) for 6 days. DMSO was used at the same concentration as vehicle control (0.1 %).

Detailed methods regarding the synthesis of the KDOBA67 compound, IC50 determination of GSK-J1 analogs, structure determination of KDM6C in complex with KDOBA67 (protein crystallisation, X-ray data collection and structure determination) are available in **SI Materials and Methods** and **Fig. S1, Table S1-S2**.

### Apoptosis and cell cycle studies, immunoblotting, qPCR, immunofluorescence

Cell cycle analysis was performed using Propidium Iodide (PI) staining, and apoptosis was determined by detecting phosphatidylserine by APC-conjugated Annexin V using flow cytometry as described in **SI Materials and Methods**. Immunoblotting and qPCRs were performed as described previously (16). Antibodies and primers are listed in **Table S3** and **Table S4** respectively. Immunofluorescence of primary chordoma cells was performed as described previously (45) using antibodies listed in **Table S3**.

### CRISPR/Cas9 knockdown

Target guide sequences validated against *KDM6A* or *KDM6B* were designed as previously published (46, 47). Guides were cloned in a CRISPR/Cas9 lentiviral backbone containing a Puromycin or a Blasticidin resistance cassette, respectively, as described previously in Shalem et al (48), and cell proliferation following genomic editing was assessed as described in **SI Materials and Methods**.

### RNA-sequencing, ChIP-sequencing, DNA methylation and data processing

Methods for chromatin immunoprecipitation (performed using a fixed ratio of *Drosophila* S2 cells ‘spiked in’ prior to fixation to allow for normalisation) and RNA extraction from cell lines, ChIPseq and RNAseq library preparation and next generation sequencing data processing are described in **SI Materials and Methods**. A list of differentially expressed genes in response to KDOBA67 (average for MUG-Chor and UM-Chor) is provided in **Dataset S2**. DNA methylation analysis was performed on formalin-fixed paraffin-embedded (FFPE) or fresh frozen tissues as described in **SI Materials and Methods**. Data are deposited in the National Center for Biotechnology Information GEO database (RNAseq and ChIPseq: GSE120214, DNA methylation: GSE119462).

## Acknowledgements

Funding for this project was received from the Chordoma Foundation (AMF, UO), Sarcoma UK (AMF, UO), Rosetrees Trust, Chordoma UK (AMF), CRUK (UO), the Oxford NIHR Biomedical Research Centre, Skeletal Cancer Action Trust (AMF), a RNOH NHS R&D grant (AMF), and the People Programme (Marie Curie Actions) of the European Union’s Seventh Framework Programme (FP7/2007–2013) (UO) under REA grant agreement n° [609305]. AMF is a National Institute for Health Research (NIHR) senior investigator. NP is a CRUK clinician scientist (Grant no:18387). LC and AMF are supported by NIHR, UCLH Biomedical Research Centre and the UCL Experimental Cancer Centre.

We thank Carina Gileadi and Kannan Velupillai for performing crystallisation experiments with KDOBA67 and KDM6C. We thank Dr. Ivana Bjedov and Victoria Martinez Miguel, UCL Cancer Institute, for providing S2 Drosophila cells, Donna Magsumbol for technical support. We thank Prof. Paolo Salomoni, Dr. Andy Feber, Teresa Sposito, Dr. Stanimir Dulev, Dr. Christopher Steele for their valuable discussion and input, UCL Cancer Institute Core Facilities, the Biobank Team at the RNOH, and the consultants who cared for the patients. We thank all the patients for donating material for this research without which it would not have been feasible.

